# Rapid High Throughput Whole Genome Sequencing of SARS-CoV-2 by using One-step RT-PCR Amplification with Integrated Microfluidic System and Next-Gen Sequencing

**DOI:** 10.1101/2020.11.04.369165

**Authors:** Tao Li, Hye Kyung Chung, Papa K. Pireku, Brett F. Beitzel, Mark A. Sanborn, Cynthia Y. Tang, Richard Hammer, Detlef Ritter, XiuFeng Wan, Irina Maljkovic Berry, Jun Hang

## Abstract

The long-lasting global COVID-19 pandemic demands timely genomic investigation of SARS-CoV-2 viruses. Here we report a simple and efficient workflow for whole genome sequencing utilizing one-step RT-PCR amplification on a microfluidic platform, followed by MiSeq amplicon sequencing. The method uses Fluidigm IFC and instruments to amplify 48 samples with 39 pairs of primers in a single step. Application of this method on RNA samples from both viral isolate and clinical specimens demonstrate robustness and efficiency of this method in obtaining the full genome sequence of SARS-CoV-2.

## INTRODUCTION

Severe acute respiratory syndrome-coronavirus 2 (SARS-CoV-2) (family *Coronaviridae*, genus *Betacoronavirus*) is responsible for the global pandemic of coronavirus disease 2019 (COVID-19) (1–4). Since its emergence in Wuhan, China, in November/December 2019, the disease has rapidly spread worldwide. As of 31 October 2020, there have been over 45 million confirmed cases, and over one million deaths in 188 countries or regions (https://coronavirus.jhu.edu/) (5–8). In addition to the continuously growing number of SARS-CoV-2 infections, increased complexity and diversity of disease symptoms are also observed. Despite the large number of whole genome sequences of SARS-CoV-2 available in GenBank and other public sources, the unprecedented scale of the viral transmission and the complexity of its pathogenic mechanism demand more genomic data to be produced using cost-effective and quality-consistent methodology (9–12).

Targeted whole genome amplification and next-generation sequencing (NGS) techniques have been used in sequencing SARS-CoV-2 genomes (8, 10, 13). In contrast to the high throughput capacity of NGS, using a conventional multiplexing reverse transcription and polymerase chain reaction (RT-PCR) procedures to amplify the large 30 Kb viral genomic RNA of SARS-CoV-2 is tedious, technically challenging, and has variable contamination risks. A rapid and streamlined approach with fewer manual steps to obtain whole genome amplicons suitable for NGS is desired.

In this study, we utilize an integrated microfluidic nucleic acid amplification system (14), custom primer design, and a one-step RT-PCR program to amplify whole SARS-CoV-2 genomes. After NGS of the amplicons, an in-house developed bioinformatics pipeline is used to rapidly obtain genome sequences with graphic summaries for data and results visualization.

## MATERIALS AND METHODS

### SARS-CoV-2 RNA samples and quantitative RT-PCR

SARS-CoV-2 RNA samples used in this study include RNA extracted from viral isolate R4717 in the US Army Medical Research Institute of Infectious Disease (USAMRIID) and RNA extracted from de-identified clinical respiratory specimens. A one-step RT-qPCR method targeting RNA dependent RNA polymerase (RdRp) was used to quantify SARS-CoV-2 RNA. Serially diluted *in vitro* transcripts (IVT), corresponding to nucleotide region 15431 to 15530 of NC_045512.2 for strain Wuhan-Hu-1, were prepared and used as real-time RT-PCR standards for quantification of genome equivalent copy number (GE) of SARS-CoV-2 RNA. Real-time RT-PCR was performed using the protocol from Carman et al. (15) with SuperScript III one-step RT-PCR System and Platinum Taq Polymerase. The QuantStudio 7 Flex Real-Time PCR System and software (ThermoFisher Scientific, Inc.) was used for data acquisition.

### Whole genome RT-PCR amplification and Illumina sequencing

For whole genome RT-PCR amplification, 35 primer pairs covering the 29903 bp SARS-CoV-2 reference genome (NC_045512.2) were custom designed by Fluidigm Corporation (South San Francisco, CA) (Table 1). The RT-PCR products are approximately 1 kb for all amplicons. One-step RT-PCR amplification was performed using Fluidigm Access Array (AA) nucleic acids amplification system (Fluidigm Corporation, CA) and SuperScript III one-step RT-qPCR System with Platinum Taq High Fidelity (ThermoFisher Scientific, Inc.). Four additional pairs of primers selected from the ARTIC network protocol v3 (https://www.protocols.io/view/ncov-2019-sequencing-protocol-v3-locost-bh42j8ye).

**Table 1.**
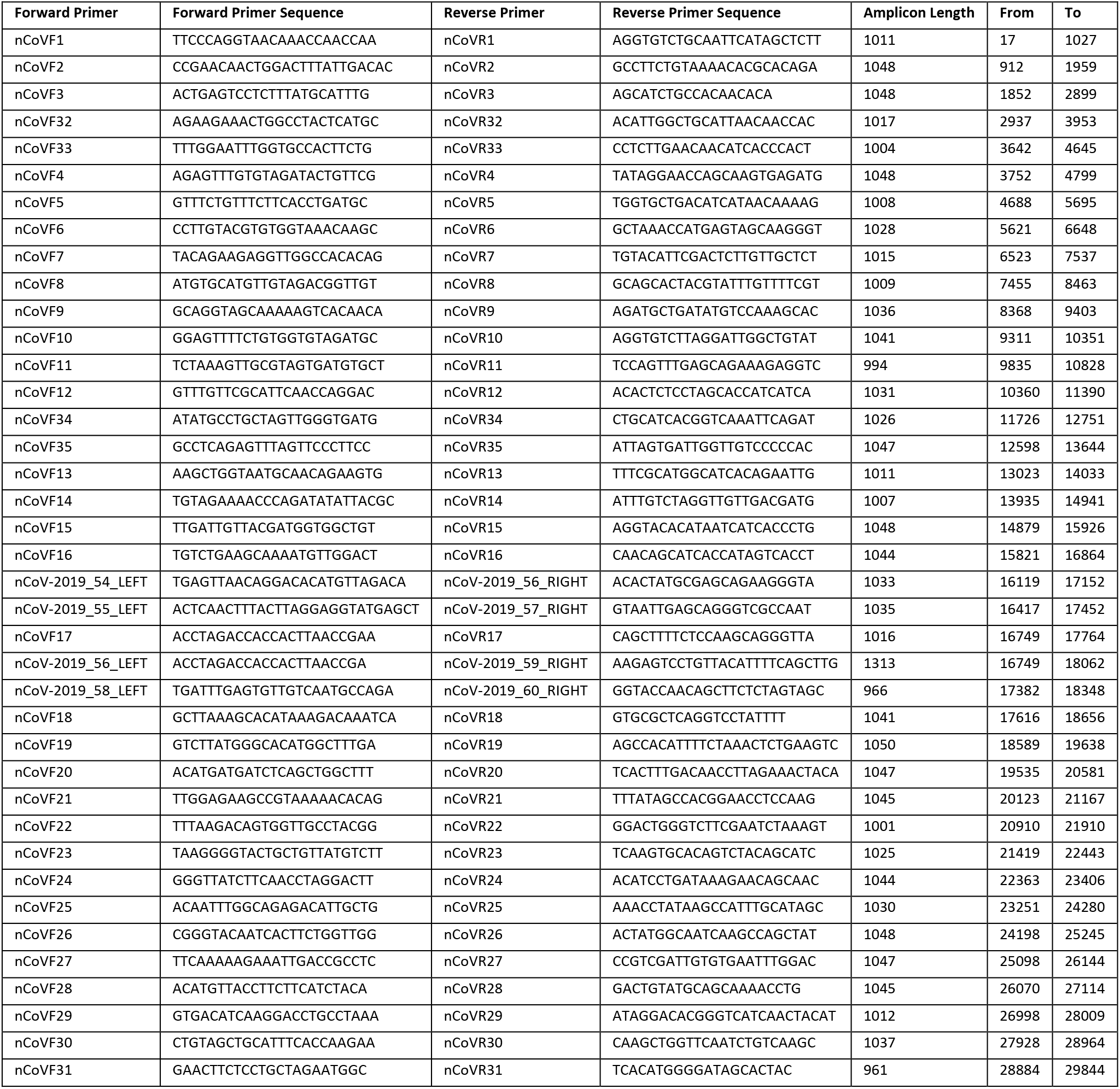
Primers for RT-PCR amplification of SARS-CoV-2 genome using Fluidigm system

SARS-CoV-2 R4717 RNA was serially diluted to the concentrations of 10^6^, 10^5^, 10^4^, 10^3^, 10^2^, 10, and 1 GE/μl (genomic equivalence per microliter). For Fluidigm RT-PCR amplification, each sample well in the Integrated Fluidic Circuit (IFC) chip contained 1.45 μl of RNA sample and 4 μl of sample mixture solution, consisting of 3 μl of 2× Reaction Mix, 0.2 μl of DMSO, 0.05 μl of RNaseOUT, 0.05 μl of SuperScript III RT/Platinum Taq High Fidelity Enzyme Mix, 0.25 μl of 20 × AA Loading Reagent, and 0.7 μl of H2O. Each well of primer mix contained 6 μl of mixture containing one pair of primers at the final concentration of 4 μM for each primer and 2.5 mM MgSO4 in AA Loading Reagent. The primed IFC chip was loaded into Fluidigm FC1 cycler and amplified using the cycling conditions: 50°C for 30 mins (reverse transcription), 94°C for 2 mins, 35 cycles of 94°C for 30 secs, 53°C for 30 secs and 68°C for 90 secs, final extension at 68°C for 7 mins, and kept at 4°C. The RT-PCR products were purified using Agencourt AMPure XP beads (Beckman Coulter, CA, USA) and then analyzed by using the Agilent Tapestation 4200 System and High Sensitivity DNA D5000 kit (Agilent Technologies, CA, USA) to determine quality and quantity of the amplicons.

The NGS libraries were prepared using Illumina DNA Flex Library kit (Illumina, CA, USA). DNA was fragmentized by tagmentation for 15 mins followed by indexing and library amplification, with 6 cycles used for 25 ng of amplicon or 12 cycles used for 1-9 ng of amplicon. The libraries were quantified by using Agilent Tapestation and DNA D5000 kit and then pooled with equal molar ratio for each library. Pooled libraries were denatured and diluted to a final concentration of 13.5 pM then sequenced using Illumina MiSeq System and Reagent Kit v3 (600 cycles).

### Reference genome mapping assembly using ngsmapper program

The full genome of SARS-CoV-2 reference NC_045512.2 was downloaded from GenBank (16) and utilized for reference-based genome assembly using the WRAIR viral disease branch in-house bioinformatics pipeline ngs_mapper version 1.5.0 (https://github.com/VDBWRAIR/ngs_mapper). This pipeline incorporates a series of quality control and assembly processes for the fastq sequence reads retrieved from the Illumina MiSeq and other platforms. These processes includes filtration to drop poorly indexed reads, read trimming based on quality thresholds using Trimommatic (17), read mapping to the reference genome using the Burrows-Wheeler Aligner with maximal exact matches (BWA-MEM) (18), read tagging, variant calling file (VCF) generation using an in-house base caller, read mapping visualizations, fastq statistic generation, and consensus sequence generation from the VCF using an in-house script basecaller.py. The pipeline streamlines bioinformatic analysis by combining multiple tools and consolidating output required for data validation and sequence curation. A minimum Phred base quality score of 35 and a minimum depth of coverage of 10 were utilized as the configuration parameters for this project. To identify variants present at a frequency of 20% or higher, an 80/20 ambiguous position threshold was used. The assembled genomes were further manually curated utilizing bam, consensus and variant calling files generated from the ngs_mapper pipeline. Geneious R10 software, integrated genome viewer (IGV) (19), and MEGA version 7 (20) were used for quality control of ambiguous calls, insertions, deletions and primer-induced mutations. The genomes were processed to only include coding sequence regions by clipping the 5’ and 3’ untranslated regions. MEGA7 was used to align genomes using default parameters with Multiple Sequence Comparison by Log Expectation (MUSCLE) (21).

## RESULTS

### Correlation of Fluidigm RT-PCR yield with amount of input SARS-CoV-2 RNA

Fluidigm RT-PCR whole-genome amplification was done in the Access Array microfluidic chip using only 1.45 μl of RNA input for each sample and with a maximum capacity of 48 samples and 48 pairs of primers. In this study, a set of 35 primer pairs, i.e. nCOVF/R1-35 in Table 1, were designed and tested for genome RT-PCR amplification by using serial dilutions of purified RNA from SARS-CoV-2 isolate R4717. The experiment was done with four replicates processed in parallel. A single band of amplicons with expected sizes of approximately 1 kb was seen for all concentrations, with the band intensities correlated with copy numbers of SARS-CoV-2 in the serial dilutions. The correlation between the concentrations of yielded amplicons and the input SARS-CoV-2 genome copy numbers (Figure 1) was significant with p-value of 3.38E-03.

**Figure 1.**
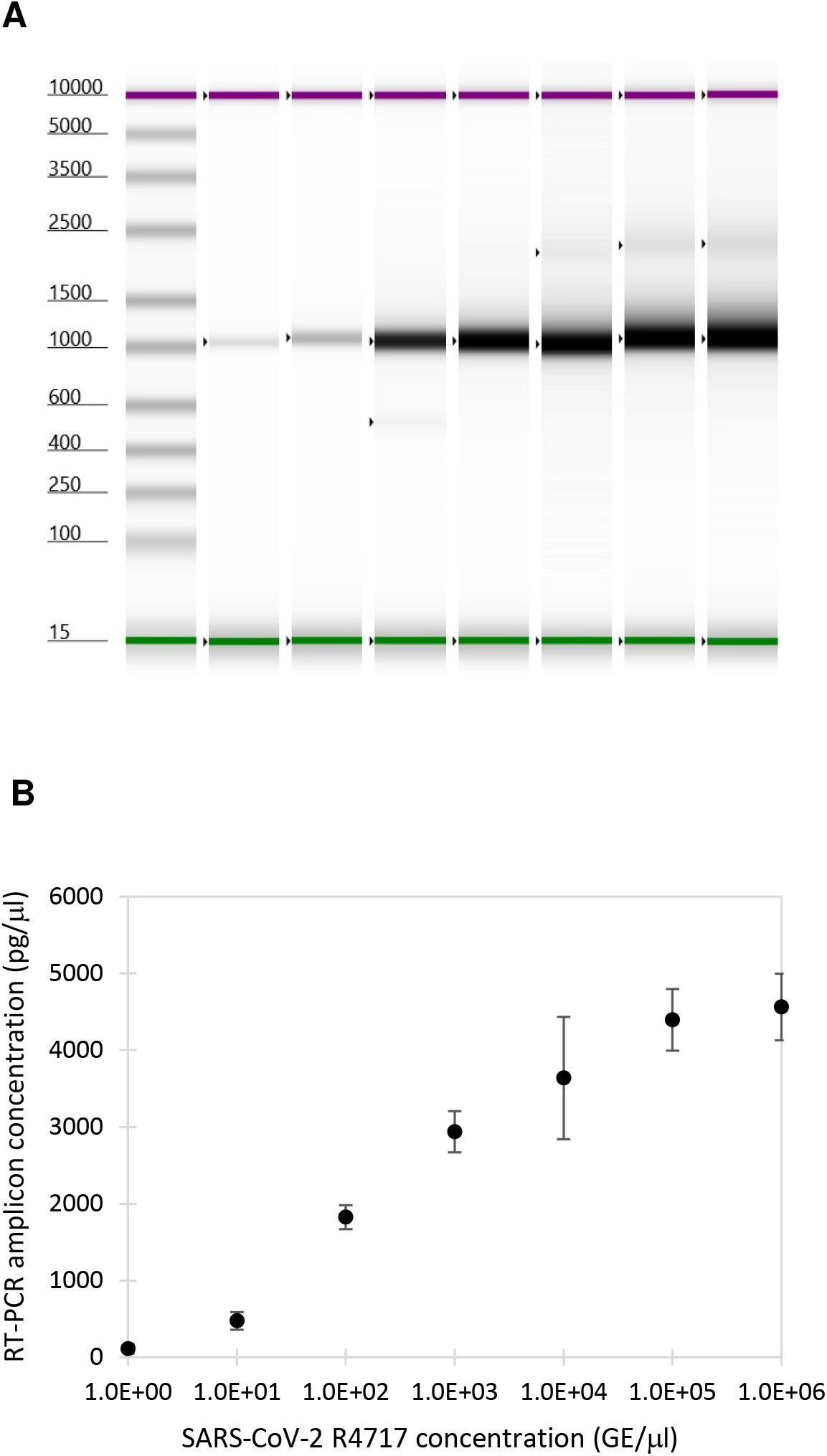
Agilent TapeStation analysis of SARS-CoV2 whole genome RT-PCR amplification. **A.** TapeStation profile of RT-PCR products. From *left* to *right*, 1, 10, 10^2^, 10^3^, 10^4^, 10^5^, and 10^6^ copies of SARS-CoV2 RNA from isolate R4717 were used. **B.** SARS-CoV-2 amplicon concentrations for RT-PCR with serial dilutions of R4717 RNA as inputs. The concentration values were the average and standard deviation from four replicates.

### Whole genome coverage and alignment depth of SARS-CoV-2 sequence assembly

MiSeq data for the quadruplicated R4717 RNA serial dilutions described above were assembled using ngs_mapper pipeline with SARS-CoV-2 complete genome sequence NC_045512.2 as the mapping reference. As expected, genome assembly results correlated well with SARS-CoV-2 copy numbers in each sample. For reactions with 10^4^ or higher SARS-CoV-2 RNA copies, complete genome sequences were readily obtained, and importantly, they had uniform coverage depth across the genome with the peaks matching the regions of amplicon alignment (Figure 2). For reactions with 10^3^ copies of SARS-CoV-2 RNA, the assembled genome sequences were nearly complete, except for two small dip/gap at positions 1870-2500 and 16800-17700. The dip/gap regions were successfully filled by adding two extra primer pairs for each region to the panel of 35 primer pairs. These additional primers were selected from the ARTIC v3 protocol, paired to cover the two regions and added into four separate Fluidigm primer wells. In total 39 pairs of primers were applied for whole genome amplification in a single RT-PCR reaction using Fluidigm nucleic acids amplification system.

**Figure 2.**
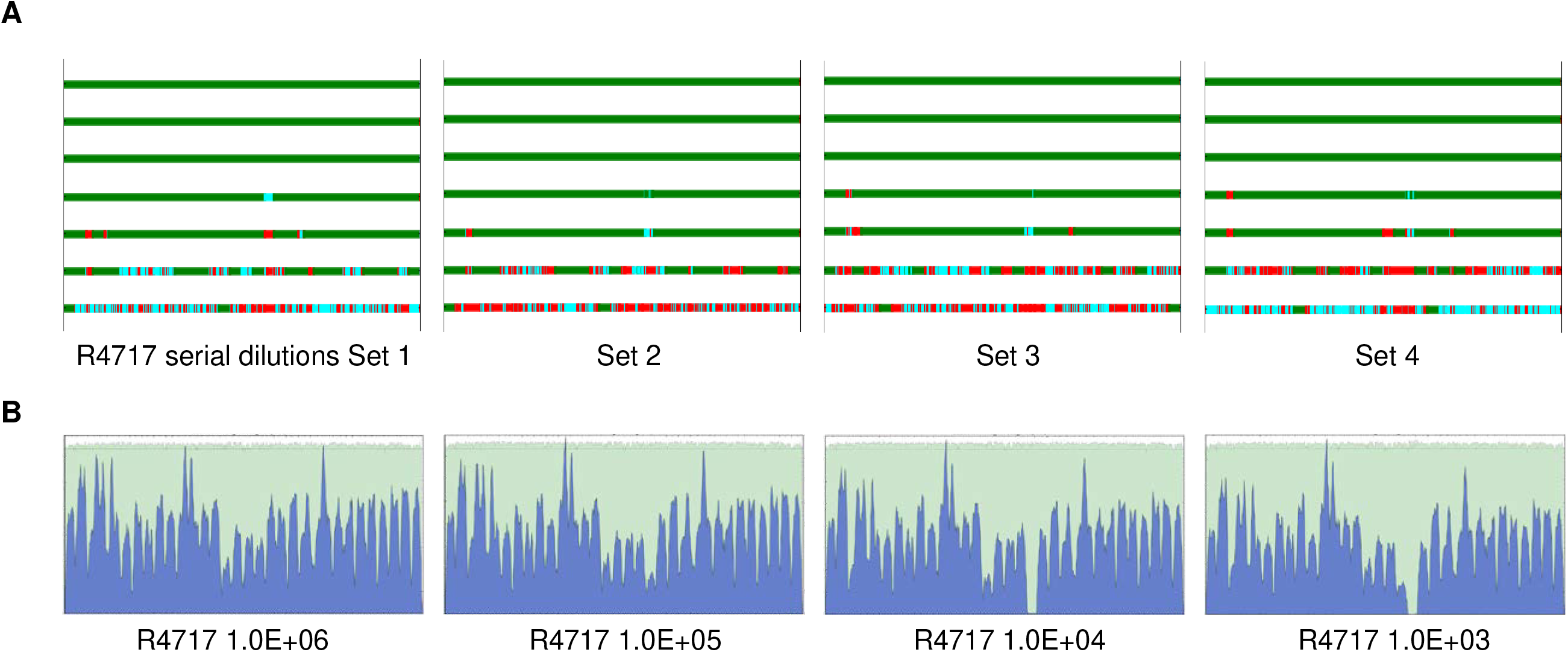
Genome coverage and sequence mapping alignment depth for MiSeq reads data. SARS-CoV-2 isolate R4717 RNA serial dilutions were subjected to Fluidigm RT-PCR amplification and MiSeq sequencing. Data were mapped using NC_045512.2 as reference. **A.** Genome coverage for 10× serial dilutions of R4717 RNA from 1.0E+06 (*top*) to 1 (*bottom*) copies. Mapping depth was indicated in colors, *green* for normal (depth ≥ 10), *cyan* for dip (depth 1-10), *red* for gap. Results for four replicates were shown. **B.** Sequence mapping graphs for 10^6^, 10^5^, 10^4^ and 10^3^ copies of RNA from one set of serial dilutions.

### SARS-CoV-2 genome sequencing of clinical respiratory specimens

This method was subsequently used in genome sequencing of SARS-CoV-2 in RNA extracts purified from COVID-19 positive nasopharyngeal swabs. The set of 29 samples contained a wide range of titers, with RT-qPCR Ct values from 13.5 to 33.4 (379 to 2.72 × 10^8^ GE/μl) of SARS-CoV-2. The RT-PCR cDNA yield has significant correlation with viral titers in the Ct range of 20 to 35, with p-value of 1.26E-05. For the samples with Ct values below 20, or exceedingly high concentrations of 1.0 × 10^7^ or greater, RT-PCR yields were substantially lower than projected (Figure 3). Importantly, this observation suggests that severe suppression of PCR could occur when samples of extremely high titer are used. Nevertheless, full or nearly complete genome coverage was obtained for all clinical specimens with titers above 1.0 × 10^4^ GE/μl, or approximate Ct value of 29.

**Figure 3.**
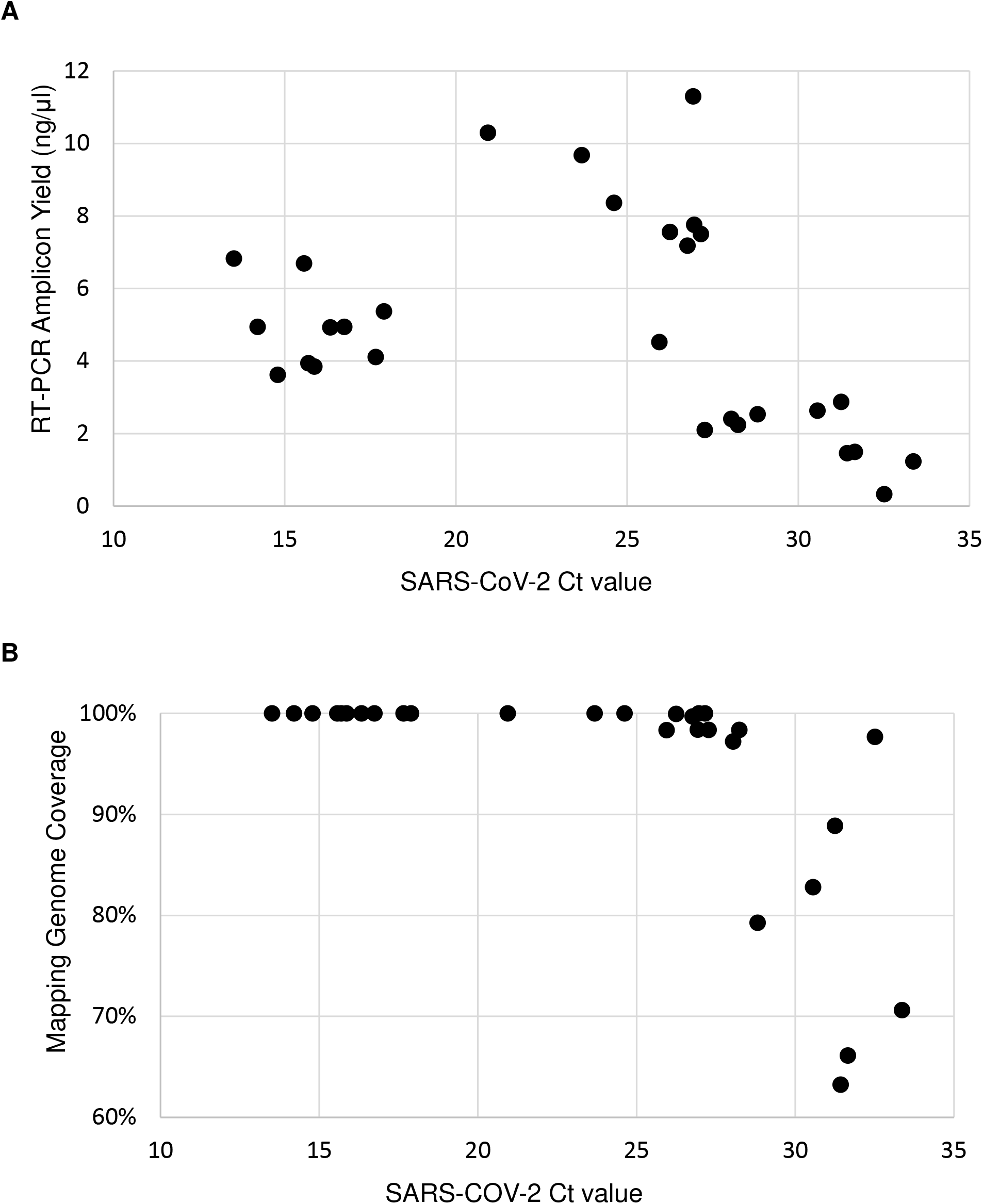
SARS-CoV2 whole genome RT-PCR amplification. Twenty nine RNA extracts from COVID-19 positive nasopharyngeal swabs were amplified and sequenced. RT-PCR yield was quantified by using Agilent TapeStation. Reference mapping coverage was determined by mapping MiSeq read data to reference genome sequence NC_045512.2.

### Comparison of assembled consensus sequences and nucleotide variations

Fluidigm one-step RT-PCR protocol was applied to SARS-CoV-2 RNA of highly varied titers. For SARS-CoV-2 isolate R4717 RNA 10× dilutions (Figure 1), a total of 14 full genomes were assembled and curated. All the full genome consensus sequences were identical except for two replicates of 1.0 × 10^4^ GE/μl. One sample had the ambiguous call Y (T or C) in the alignment nucleotide position 2105, while the other three replicates had a C. Another sample had a T in position 23260 while all the other three had the ambiguous call Y. These were the only ambiguous positions found in the samples, showing low sample diversity at a variant frequency of 20% or higher. Together this sequencing approach produces high accuracy results.

## DISCUSSION

Even before the disease was named as COVID-19, the sequence was swiftly determined using next-gen sequencing technologies and SARS-CoV-2 was identified as the causative pathogen for the emerging acute respiratory disease (5, 22). The first sequence was made publicly available immediately with a massive number of sequences subsequently generated and shared, which has greatly facilitated research and development (23–25). All the efforts in developing, improving, and sharing materials and/or methods has played an essential role in sequence-based investigations. All the known sequencing protocols use conventional PCR apparatuses with differences in design and selection of primers and reaction parameters. In this study, we applied one-step RT-PCR protocol on a microfluidic platform (14) to establish a convenient workflow with throughput, speed, simplicity, consistency, and yield suitable for COVID-19 genome sequencing. Fluidigm Access Array IFC holds 48 RNA samples (inlets) and 48 primer pairs (inlets). Steps for mixing sample with primers are obviated, which not only substantially reduces pipetting manipulation but also effectively mitigates the chance of sample-to-sample cross-contamination. In this report, we selected 39 primer pairs to obtain even genome coverage. Nine more individual pairs of primers can be easily added to the panel of 48 primer inlets to quickly address emerging SARS-CoV-2 genetic divergence. Moreover, the total number of primer pairs can be further increased without difficulty by pooling together several compatible primer pairs and adding them into one primer inlet. Each sample is mixed with individual primer pairs in IFC microfluidic chambers for nano-liter (nl) RT-PCR amplifications in a simplex independent reaction manner. In contrast, conventional PCR methods often need optimization of primer pooling and reaction parameters to circumvent primer-to-primer interference and to avoid highly variable yields among amplicons.

Since input RNA samples are partitioned into individual nl reaction chambers to cross-mix with individual primer pairs, microfluidic applications including Fluidigm IFC require sufficient genomic copies in order to achieve whole genome amplification. In consequence, Fluidigm amplification based whole genome sequencing has limitations in sequencing low titer samples. For RNA samples with 1000 GE/μl or lower concentration, multiplex RT-PCR based methods or SARS-CoV-2 hybridization-based enrichment method might be a more suitable choice for whole genome sequencing (26–28). The addition of a first-strand cDNA synthesis step prior to Fluidigm amplification, with a change of Fluidigm thermocycling program from RT-PCR to PCR, may help increase genome coverage for low titer samples. Many, if not all, COVID-19 specimens are tested with quantitative molecular tests and the Ct values or equivalent titer scores are readily available for deciding whether one-step Fluidigm amplification-based genome sequencing protocol is appropriate.

Using this method, very few sequence assembly errors were observed throughout the tested SARS-CoV-2 sample genomes. These errors might be due to PCR, sequencing or basecalling algorithm errors, as well as due to normal fluctuations in the minor variant frequencies between the sample aliquots. When assembled sequences show potentially significant nucleotide alterations or indels, thorough examination of the data quality processing, primer trimming, curation of sequence assembly, and detailed laboratory records are needed. Whenever possible, running replicate samples, repeating the experiment entirely, or using other methods are important to validate genome variations and minor variants.

In conclusion, our study demonstrates a convenient SARS-CoV-2 whole genome sequencing protocol by incorporating one-step RT-PCR amplification, microfluidic technology, and next-generation sequencing to achieve a simple and fast workflow with consistent and quality data. The performance of the protocol was verified using viral isolate RNA and tested by sequencing clinical respiratory samples of varying viral titers.

## Acknowledgments

We thank Mr. James S. Hilaire, Ms. Nicole R. Nicholas, Mr. Tuan K. Nguyen and Ms. April N. Griggs for their assistance in project management, sample tracking, storage and retrieval.

## Disclaimer

Material has been reviewed by the authors’ respective institutions. There is no objection to its presentation and/or publication. The views expressed here are those of the authors and do not reflect the official policy of the Department of the Army, Department of the Navy, Department of Defense or U.S. Government. This is the work of U.S. government employees and may not be copyrighted (17 USC 105).

